# The noncoding RNA *PRANCR* regulates splicing of *Fibronectin-1* to control keratinocyte proliferation and migration

**DOI:** 10.1101/2021.06.22.449364

**Authors:** Auke B.C. Otten, Oyumergen Amarbayar, Pengfei Cai, Binbin Cheng, Kun Qu, Bryan K. Sun

## Abstract

Most human genes undergo alternative splicing (AS), but the regulation and functional consequences of most splicing events remain unknown. Long noncoding RNAs (lncRNAs) have recently been discovered to have novel roles in the regulation of AS. Here we investigate whether *PRANCR*, a lncRNA recently identified to be essential for epidermis formation, functions by controlling AS of cell fate genes. Using transcriptome-wide analysis, we identified 238 exonic splicing events regulated by *PRANCR*. Among these is alternative splicing of an exon containing the extra domain A (EDA) in the gene fibronectin-1 (*FN1*). Expression of the FN1-EDA+ isoform is enriched in proliferating keratinocytes. We find that *PRANCR* regulates EDA inclusion by controlling expression of the serine/arginine-rich splicing factors (SRSFs) 1 and 7. Depletion of *PRANCR* or FN1-EDA resulted in decreased proliferation, increased CDKN1A/p21, and inhibition of keratinocyte migration. We find that these cellular phenotypes can be explained by reduced phosphorylation of focal adhesion kinase (FAK). Collectively, these results identify a lncRNA regulating skin function through alternative splicing of a cell fate gene.

## Introduction

The development of deep sequencing techniques (Pan *et al*, 2008; Wang *et al*, 2008) and bioinformatic pipelines (Shen *et al*, 2014; Sterne-Weiler *et al*, 2018) have enabled researchers to comprehensively map changes in alternative splicing (AS) of RNA transcripts during cell differentiation, tissue development and disease (Scotti & Swanson, 2016). AS enables the generation of multiple mRNA isoforms from the same gene, which can diversify their cellular functions and characteristics. Currently, over 100,000 splicing events have been identified, occurring in ~95% of all human genes (Pan *et al*, 2008; Wang *et al*, 2008). Though AS events are ubiquitous, the molecular mechanisms governing AS are incompletely understood, as are the functional consequences of most splicing events (Baralle & Giudice, 2017).

Splicing is catalyzed by the spliceosome, a ribonucleoprotein complex consisting of small nuclear RNAs and RNA binding proteins (RBPs) that function as splicing factors (SFs) (Chen & Cheng, 2012). Regulation of splicing can occur by control of SF expression, activity, and/or cellular localization (Chen & Cheng, 2012). Emerging evidence demonstrates roles for long noncoding RNAs (lncRNAs), non-protein coding RNA transcripts >200 nucleotides in length, in AS regulation (Romero-Barrios *et al*, 2018). LncRNAs can modify the expression of specific SFs to control AS events (He *et al*, 2019). Alternatively, some lncRNAs regulate posttranslational modifications of SFs or function as scaffolds or ‘hijackers’, affecting the ability of SFs to interact with RBPs or mRNAs (Romero-Barrios *et al*, 2018). One well-characterized example is the lncRNA *MALAT1*, which can modulate the expression of the SF *RBFOX2* (Gordon *et al*, 2019), alter the phosphorylation status and abundance of serine/arginine-rich SF1 (SRSF1) (Tripathi *et al*, 2010), and prevent formation of a splicing regulatory complex by hijacking SFPQ (Ji *et al*, 2014). While *MALAT1* demonstrates the proof of principle that lncRNAs can regulate AS, it is still unclear if this is a prevalent mechanism by which lncRNAs function (Romero-Barrios *et al*, 2018; Statello *et al*, 2021).

Recently, we identified *PRANCR* as a novel lncRNA required for epidermal homeostasis (Cai *et al*, 2020). The epidermis is a stratified epithelium that serves as the protective outermost layer of the skin. It undergoes continuous renewal, requiring a balance between keratinocyte proliferation and differentiation (Fuchs & Raghavan, 2002). *PRANCR* depletion causes epidermal keratinocytes to lose both their proliferative and differentiative capacity and inhibits the formation of the epidermal barrier in organoid models. In part, *PRANCR* functions by controlling expression of late cell cycle genes, including CDKN1A/p21 (Cai *et al*, 2020). However, depletion of *PRANCR* did not change the steady state mRNA expression of canonical keratinocyte cell fate (KCF) genes (Wu *et al*, 2012), indicating that it may function through non-transcriptional mechanisms as well.

Keratinocyte behavior can be controlled by expression of specific splicing isoforms of regulatory genes (Tanis *et al*, 2018). For instance, specific mRNA isoforms of dermokin (*DMKN*), desmoplakin (*DSP*), fibroblast growth factor receptor 2 (*FGFR2*) and *TP63* are expressed in epidermal keratinocytes in either the progenitor or differentiated states (Naso *et al*, 2007; Cabral *et al*, 2012; Ranieri *et al*, 2016; Warzecha *et al*, 2009; Yang *et al*, 1998). However, the repertoire of alternative mRNA splicing through the biological process of epidermal differentiation has not been fully characterized. Here, we investigated if *PRANCR* functions by regulating splicing of KCF genes. We demonstrate that *PRANCR* controls splicing of the exon containing extra domain-A (EDA) in the KCF gene fibronectin-1 (*FN1*) and define a mechanistic role for *PRANCR* and FN1-EDA during keratinocyte proliferation and migration. These findings bring new insight to the phenotype of impaired wound reepithelization previously observed in *FN1*-EDA knockout mice and implicate a role for lncRNAs in keratinocyte mRNA splicing regulation.

## Materials and Methods

### Cell culture

Primary epidermal keratinocytes were isolated from discarded neonatal foreskins generated from elective circumcisions, collected with written informed consent under an Institutional Review Board protocol approved by the University of California, San Diego. Cells were isolated as described previously (Aasen & Belmonte, 2010) and propagated in a 50:50 mixture (“50:50 media”) of K-SFM and 154 media (Life Technologies) with recommended supplements and 1x Antibiotic-Antimycotic (Thermo Fisher Scientific), at 37°C and 5% CO_2_. To induce keratinocyte differentiation, primary epidermal keratinocytes were seeded at confluency in the presence of 1.2 mM CaCl_2_ for four days. For experiments with TGF-β1, recombinant human TGF-β1 (Life Technologies) was prepared in sterile water with 0.1% BSA following the manufacturer’s instructions and diluted in 50:50 media.

### Alternative splicing events analysis

RNA-sequencing of normal and *PRANCR*-depleted keratinocytes (n=4 each, NCBI Gene Expression Omnibus (GEO) accession GSE125400 (Cai *et al*, 2020)) was assessed with rMATs (version 3.2.5) to identify differential exon usage between control and *PRANCR* knockdown cells. Significant AS events were defined as exons displaying a difference in the <u>p</u>ercentage <u>s</u>pliced <u>i</u>n (PSI) between control and knockdown of at least 0.1 and having a false discovery rate (FDR) below 0.05.

### RNA isolation, RT-PCR and qPCR

RNA was prepared by collecting cells in TRIzol reagent and isolated using the Direct-zol RNA Kit (Zymo Research) following manufacturer’s recommendations. RNA was reverse transcribed using the iScript cDNA synthesis Kit (Bio-Rad). For determination of the *FN1*-EDA isoform ratio, cDNA was amplified with DreamTaq Green PCR Master Mix (ThermoFisher) and visualized on a 1% agarose gel in the presence of SYBR Safe (Invitrogen). Relative quantification of *FN1*-EDA+ and *FN1*-EDA- bands was performed with Image Lab Software (Bio-Rad). Primers used (5’ to 3’): *FN1*-Fwd, TATTGAAGGCTTGCAGCCCA, *FN1*-Rev, GAGCTGTCAGGAGCAAGGTT.

For qRT-PCR, PCR amplification reactions were performed using iTaq Universal SYBR Green Supermix (Bio-Rad) on a CFX96 Touch Real-Time PCR Detection System (Bio-Rad). Gene expression analysis was performed using the 2^−ΔΔCT^ method. Primers (5’-to-3’): SRSF1-Fwd, GCGGTCTGAAAACAGAGTGG, SRSF1-Rev, TGCCATCTCGGTAAACATCA, SRSF3-Fwd, GCAGTCCGAGAGCTAGATGG, SRSF3-Rev, TGGGCCACGATTTCTACTTC, SRSF7-Fwd, TGCAGTACGAGGACTGGATG, SRSF7-Rev, GGGCAGGTGGTCTATCAAAA, FN1-Fwd, AGCGGACCTACCTAGGCAAT, FN1-Rev, TCACCCACTCGGTAAGTGTTC, L32-Fwd, AGGCATTGACAACAGGGTTC, L32-Rev, GTTGCACATCAGCAGCACTT.

### RNA interference-mediated gene knockdown and overexpression experiments

For short hairpin (sh)-targeted gene knockdown of *PRANCR*, *SRSF1, 3* and *7*, as well as specific *FN1*-isoforms, shRNAs were cloned into the pLKO.1 vector (the RNAi consortium). shRNA sequences (5’-to-3’): shSCR-1, CCTAAGGTTAAGTCGCCCTCG, shSCR-2, GCAAGCTGACCCTGAAGTTCA, shSRSF1-1, GTGGAAGTTGGCAGGATTTAA, shSRSF1-2, GTTTGTACGGAAAGAAGATAT, shSRSF3-1, GCTAGATGGAAGAACACTATG, shSRSF3-2, GAAGTGGTGTACAGGAAATTA, shSRSF7-1, ATCTAGGTCTGGTTCTATAAA, shSRSF7-2, GACTGGATGGAAAGGTGATTT, shFN1-3’UTR-1, GATCTTGTTACTGTGATATTT, shFN1-3’UTR-2, ACCTTTCCTTCATTGAATAAA, shFN1-EDA-1, GGTTGCCTTGCACGATGATAT, shFN1-EDA-2, GGATGGAATCCATGAGCTATT, shPRANCR-1, CACTTTGAATGACAACGATTT, shPRANCR-2, GCAAGCTGACCCTGAAGTTCA, shPRANCR-3, TTCCACCCAAGCCACAATAAT.

For overexpression experiments, open reading frames were PCR-amplified using CloneAmp HiFi PCR Premix (Clontech), gel-purified using the Nucleospin^®^ Gel and PCR Clean-Up kit (Macherey-Nagel), and cloned into the LentiORF pLEX-MCS Vector (Thermo Scientific) using In-Fusion cloning (Clontech). Primers used: SRSF1-F, CTCTACTAGAGGATCCATGTCGGGAGGTGGTGTG, SRSF1-Rev, TAGACTCGAGCGGCCGCTTATGTACGAGAGCGAGATCTGC, SRSF3-Fwd, CTCTACTAGAGGATCATGCATCGTGATTCCTGTCC, SRSF3-Rev, TAGACTCGAGCGGCCCTATTTCCTTTCATTTGACCTAGAT, SRSF7-Fwd, CTCTACTAGAGGATCCATGTCGCGTTACGGGCGG, SRSF7-Rev, TAGACTCGAGCGGCCGCTCAGTCCATTCTTTCAGGACTTGC.

Lentivirus was generated by transfection of packaging and transfer plasmids into 293T cells using polyethylenimine. Lentiviral particles were collected 72 h after transfection and concentrated with Lenti-X Concentrator (Clontech) and stored at − 80°C. For knockdown experiments, up to 5.0×10^5^ keratinocytes were infected in 50:50 medium containing 3 μg/mL polybrene and incubated overnight. Infected cells were selected in medium supplemented with 1 μg/mL puromycin for 72 hours.

### Cell proliferation assay

To assess cell proliferation rates, 5,000 cells were seeded on a 24-well plate in duplicate for each condition and each time point. Media was changed every 48 hours. At each time point, viable cell abundance was assessed using resazurin (alamarBlue; Thermo Fisher Scientific). Fluorescence was measured after 2 hours of incubation at 37°C using the SpectraMax® iD3 microplate reader (Molecular Devices). To compare growth rates, fluorescence intensity at the start of the experiment (day 0) was set to 1. Proliferation was assessed relative to the starting value. Cell cultures remained sub-confluent in all conditions for the duration of the experiment.

### Protein isolation and western blotting

Whole-cell protein lysates were collected in IP Lysis Buffer (Pierce) and quantitated with a BCA assay (Pierce). Equal amounts of protein lysates were loaded onto 4-12% Bis-Tris gels (Invitrogen) and resolved by electrophoresis using Bolt® MOPS SDS Running buffer (Invitrogen). Wet transfer was performed onto PVDF membranes, and primary antibodies were incubated at 4°C overnight. Membranes were washed 3 times in PBS-0.1% Tween, then incubated with fluorescent secondary antibodies according to the manufacturer’s recommendations (Li-Cor), washed and visualized. Blots were imaged on an Odyssey® imager (Li-Cor) and densiometric quantitation was performed using Image Studio software (Li-Cor). Antibodies and dilutions used: ACTB (Rabbit, 1:1,000, Cell Signaling Technologies #4967), TUBB (Mouse, 1:1,000, Development Studies Hybridoma Bank (E7)), GAPDH (Mouse, 1:200, Santa Cruz Biotechnologies #47724), FN1-total (Mouse, 1:200, Santa Cruz Biotechnologies #11825), FN1-EDA (Mouse, 1:200, Santa Cruz Biotechnologies #59826), KRT10 (Rabbit, 1:500, Abcam #76318), CDKN1A/p21 (Rabbit, 1:1,000, Cell Signaling Technologies #2947), total-FAK (Rabbit, 1:1,000, Cell Signaling Technologies #3285) and phosphorylated FAK (Tyr397, Rabbit, 1:1,000, Cell Signaling Technologies #8556).

### Expression microarray analysis

A microarray dataset (GSE136375) assessing transcriptional changes upon calcium-induced keratinocyte differentiation was re-analyzed using GEO2R. Datasets were divided into “progenitors”, “3 days differentiation” and “6 days differentiation”. Default GEO2R settings were used to calculate false discovery rates (Benjamini & Hochberg) and annotation of the Agilent-014850 Whole Human Genome Microarray was generated using the NCBI database. From the results, SRSF gene names were retrieved and ordered according to the calculated Log2 fold change.

### *In vitro* wound healing migration assay

To assess migration capacity, 37,000 cells per well were seeded into each well of 3-well silicone inserts on tissue-culture treated surfaces (Ibidi GmbH, Germany). After confirming full confluency, inserts were removed 18 hours after seeding, resulting in two 500 μm cell-free gaps with confluent cells at both sides of the gap. Migration was assessed by imaging the gap at three fixed positions using the EVOS M5000 microscope (ThermoFisher Scientific) at different timepoints until full closure of the control condition (shSCR). To measure wound healing size, each picture was analyzed in *ImageJ* using the *Wound_healing_size_tool* plugin (Suarez-Arnedo *et al*, 2020). Cell-free areas (Area pixels^2) were normalized to the starting cell-free gap size.

### Statistical analysis and regression analysis

All data were statistically analyzed using GraphPad (Prism, La Jolla, CA). Unless stated otherwise, differences between experimental groups and control conditions (shSCR) were tested using *t*-tests and raw *p*-values are depicted. For quantification of Western Blots, band intensities were normalized to housekeeping genes (TUBB, ACTB and/or GAPDH) and compared against controls (average of shSCR-1 and shSCR-2).

## Results and Discussion

### *PRANCR* regulates alternative splicing of the cell fate gene *fibronectin-1*

We recently identified a novel role for the lncRNA *PRANCR* in epidermal proliferation and differentiation (Cai *et al*, 2020). Because stratified epidermis depends critically on the master transcription factor p63, we initially hypothesized that *PRANCR* would influence expression of p63-controlled keratinocyte cell fate (KCF) genes (Wu *et al*, 2012). However, despite the strong effect of *PRANCR* depletion on keratinocyte proliferation and differentiation, we did not identify notable changes to expression of any KCF genes (Supplemental Figure 1A).

Recently, it was described that some lncRNAs can regulate alternative splicing of mRNAs. We therefore assessed if *PRANCR* might function by controlling expression of functional mRNA isoforms. To define the repertoire of *PRANCR*-regulated splicing events, we quantitated differential mRNA isoform expression in RNA-seq data of control and *PRANCR*-depleted primary human keratinocytes (*n* = 4 each) (Cai *et al*, 2020). Using the <u>r</u>eplicate <u>m</u>ultivariate <u>a</u>nalysis of <u>t</u>ranscript <u>s</u>ilencing (rMATS) pipeline, a statistical method for detection of AS from replicate RNA-seq data (Shen *et al*, 2014), we identified 238 exons with differential abundance (FDR <0.05 and a difference in <u>p</u>ercentage <u>s</u>pliced <u>i</u>n (PSI) > 0.1) in wild-type vs. *PRANCR*-depleted keratinocytes (Figure 1A). Among these, two KCF network genes showed significant isoform expression differences: fibronectin-1 (*FN1*) and the protein tyrosine phosphatase *PTPN12* (Figure S1B-C).

**Figure 1.**
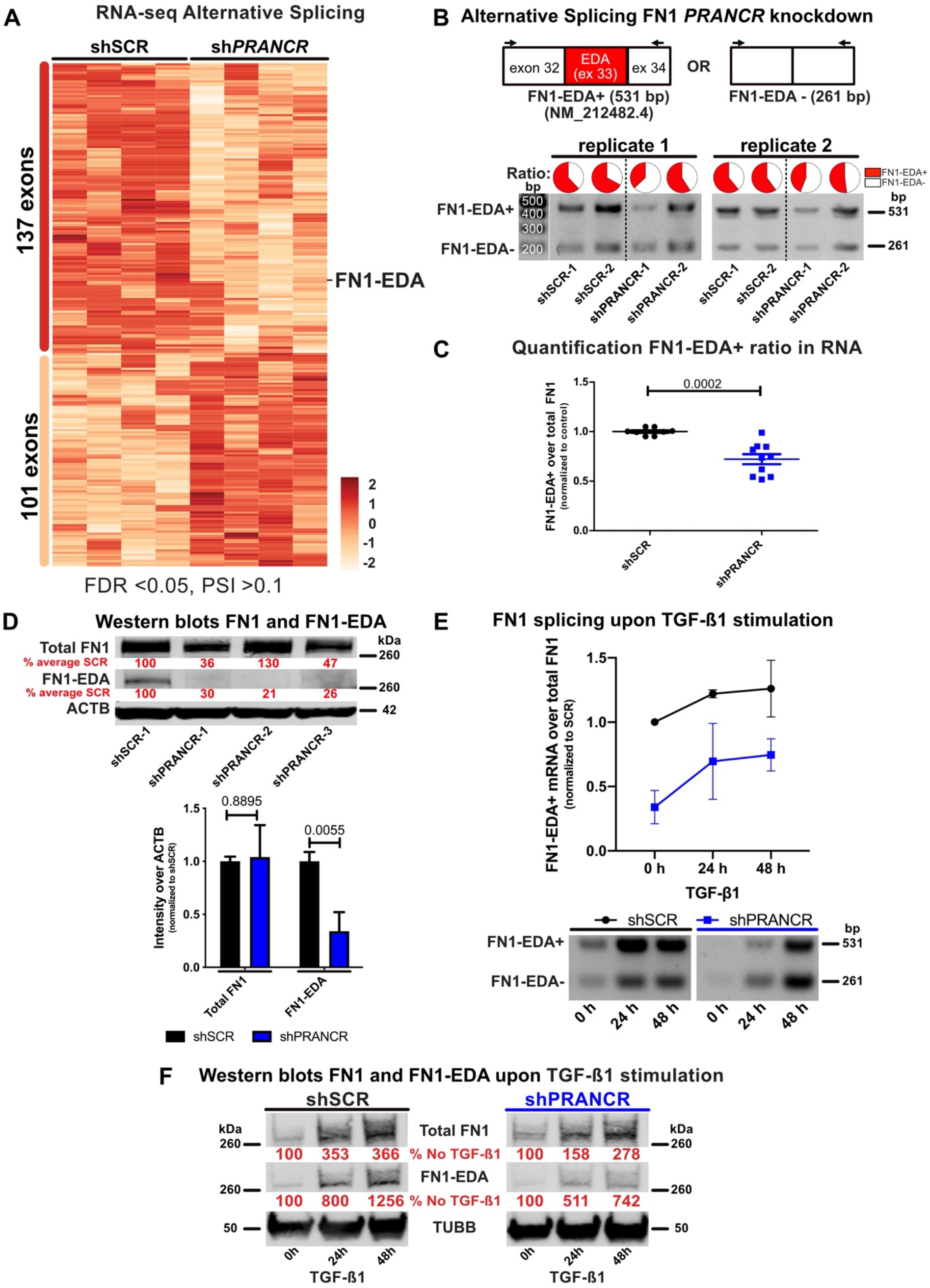
*PRANCR* regulates splicing of the fibronectin-1 EDA domain-containing isoform. A) Hierarchical clustering heat map of genes with a difference in <u>p</u>ercentage <u>s</u>pliced <u>i</u>n (PSI) >0.1 between *PRANCR*- depleted keratinocytes (shPRANCR1/2, *n* = 4) and control samples (shSCR1/2, *n* = 4), and FDR < 0.05 (Cai *et al*, 2020). B) *Top*: diagram indicating the EDA domain in the *FN1* gene and the location of primers used for RT-PCR with predicted amplification sizes. *Bottom*: analysis of AS of *FN1* mRNAs from control and *PRANCR*-depleted primary keratinocytes by RT-PCR. Representative gels are shown of two replicate experiments with ratios of *FN1*-EDA+ over total *FN1*, following band intensity quantification and adjustment for differences in amplicon size (non-adapted gel pictures depicted in supplemental Figure S1D). C) Graph showing quantification of *FN1-EDA* over total *FN1* normalized to controls. Each dot represents the outcome of an independent experiment. Bars, mean with SEM; *n* = 8 in control and *n* = 10 in *PRANCR*-knockdown (different shRNAs combined). D) Western blot of FN1 and FN1-EDA in control (shSCR) and *PRANCR*-depleted (using three independent shRNAs, sh*PRANCR*1-3) cells. Protein abundances are shown as a percentage of SCR after blot quantification. (non-adapted Western blot pictures depicted in supplemental Figure S2A). E) Analysis of AS of *FN1* mRNAs from control and *PRANCR*-depleted primary keratinocytes stimulated with 10 pmol TGF-β1. Graph shows changes in *FN1*-EDA+ over total FN1, normalized to controls at *t* = 0. *n* = 2 for each time point for each experimental condition. (non-adapted gel pictures depicted in supplemental Figure S1F). F) Western Blot of FN1- and FN1-EDA in control and *PRANCR*-depleted keratinocytes upon stimulation with 10 pmol TGF-β1. Protein abundances are shown as a percentage of t=0 after blot quantification by densitometry (non-adapted western blot pictures depicted in supplemental Figure S2C).

Fibronectin-1 regulates keratinocyte differentiation (Adams & Watt, 1989), adhesion and proliferation by binding and activating the integrin-signaling pathway (Watt, 2002). *PRANCR* depletion resulted in reduced expression of the *FN1* mRNA isoform that encodes for extra-domain A (EDA; or extra-type III homology; schematic in Figure 1B). The EDA domain has important cellular functions across cell types and is known to regulate cell cycle progression, differentiation and regulation of gene expression (Jarnagin *et al*, 1994; Xia & Culp, 1995; Manabe *et al*, 1999; Guan *et al*, 1990; Saito *et al*, 1999). *FN1*, and specifically the *FN1*-EDA isoform, are essential for murine embryonic development and full life span (Astrof *et al*, 2007; Muro *et al*, 2003). The location and timing of *FN1*-EDA inclusion are important for its final function. Mice specifically lacking *FN1*-EDA display abnormal myofibroblast differentiation during lung fibrosis (Muro *et al*, 2008; Kohan *et al*, 2011), as well as abnormal reepithelization during skin wound healing (Muro *et al*, 2003). In contrast, elevated levels of the *FN1*-EDA^+^ isoform has been identified in cancers, psoriasis, rheumatoid vasculitis and diabetes, suggesting a strong association of the EDA-domain with disease (Szell *et al*, 2004; Guban *et al*, 2016; McFadden *et al*, 2011; Voskuyl *et al*, 1998; Kanters *et al*, 2001). The observation that *PRANCR* affected *FN1-EDA* splicing in keratinocytes prompted us to investigate this splicing event and its functional consequences in greater detail.

We first reassessed *FN1* mRNA isoform expression using semi-quantitative RT-PCR and confirmed that proportional expression of EDA-containing mRNA isoforms (hereafter termed *FN1*-EDA+) decreased significantly in *PRANCR*-depleted keratinocytes compared to controls (Figures 1B-C and Supplemental Figure S1D). To assess if mRNA changes correlated to differential protein isoform expression, we performed immunoblots using two distinct FN1 antibodies, one that targets a universal epitope common to all FN1 isoforms (total FN1) and one that specifically detects the EDA domain (FN1-EDA). *PRANCR* depletion led to a reduction of FN1-EDA+ protein expression by 70-80% (Figure 1D and Supplemental Figure S2A), reflecting a strong effect of *PRANCR* on FN1 protein isoform expression.

FN1 splicing can be induced by transforming growth factor bèta-1 (TGF-β1; (Ventura *et al*, 2018) and Supplemental Figures S1E and S2B). We therefore assessed if the induction of *FN1*-EDA splicing was affected by *PRANCR*. Following TGF-β1 stimulation, *PRANCR*-depleted keratinocytes displayed decreased *FN1*-EDA+ mRNA (Figure 1E and Supplemental Figure S1F) and protein abundance (Figure 1F and Supplemental Figure S2C). Together, these results showed that *PRANCR* affects splicing of the *FN1*-EDA exon in both unperturbed keratinocytes and following TGF-β1 stimulation.

Expression of cell fate genes can regulate the transition between cell states. During homeostasis, the epidermis maintains a balance between replicating progenitors and post-mitotic, differentiated keratinocytes (Fuchs & Raghavan, 2002). We therefore evaluated the expression of *FN1*-EDA+ in progenitor vs. differentiated keratinocytes (Supplemental Figure S3A). We observed two changes in the pattern of *FN1* expression. First, there was a decrease in overall *FN1* expression in differentiated keratinocytes compared to progenitors. Second, there was a significant reduction in the relative expression of the *FN1*-EDA+ isoform (Figure 2A-B and Supplemental Figure S3B). These two patterns of *FN1* mRNA expression were consistent with what was observed on the protein level, as assessed by immunoblotting (Figure 2C and Supplemental Figure S3C). Together, these results confirmed a correlation between enriched *FN1*-EDA+ isoform expression and the replicative progenitor state. This dynamic regulation of the relative expression of *FN1*-EDA+ and EDA– isoforms is also observed during cutaneous wound healing (Ffrench-Constant *et al*, 1989) and led us to consider a potential role of *PRANCR* as a switch for regulating *FN1* isoform expression and keratinocyte cell fate.

**Figure 2.**
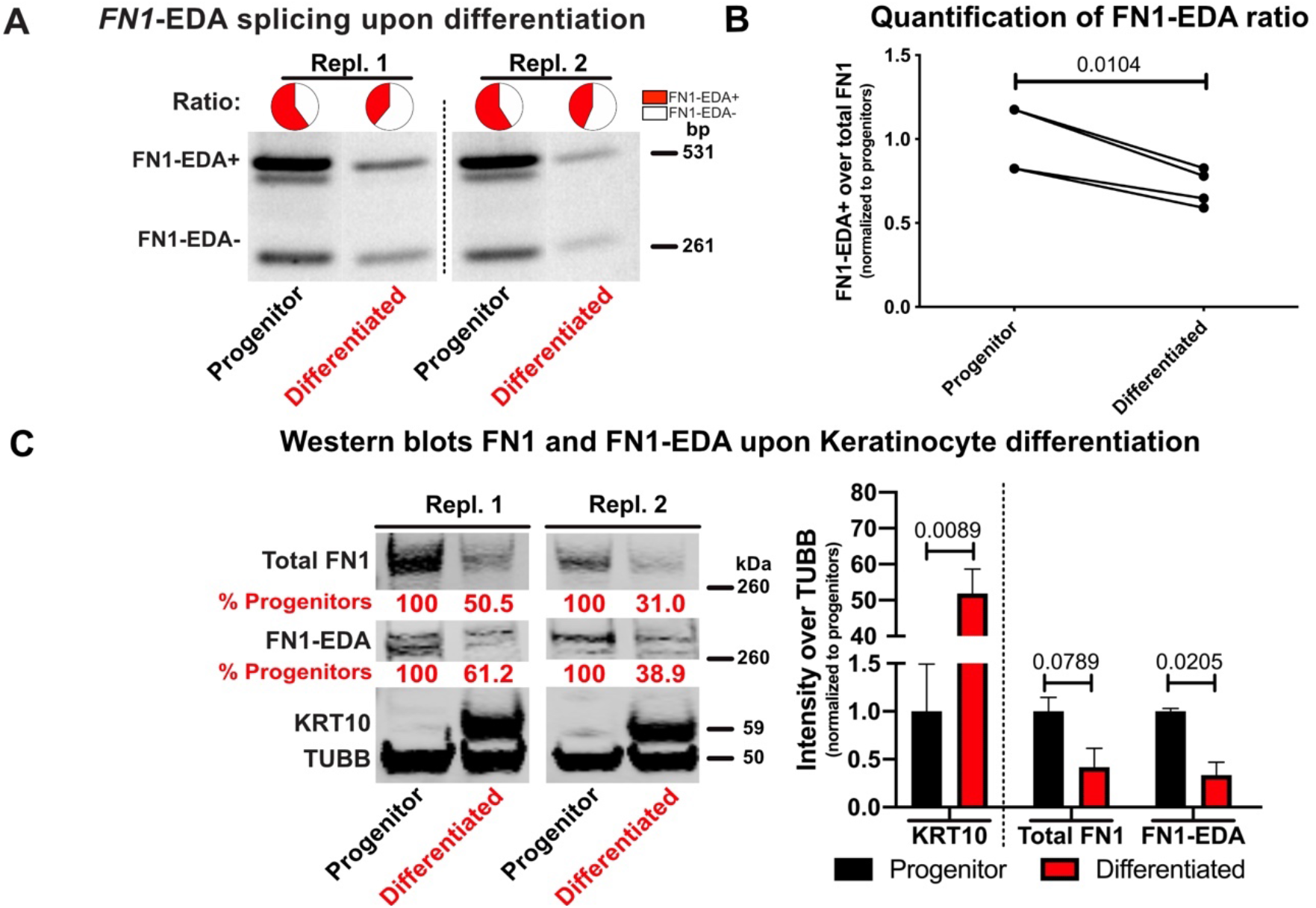
FN1-EDA splicing changes upon keratinocyte differentiation. A) Representative gels showing differences in *FN1*-EDA+ mRNA abundance upon 4 days of calcium-induced differentiation (as assessed by induction of known epidermal differentiation markers (Supplemental Figure 3A)) in two replicate experiments with ratios of *FN1*-EDA+ over total *FN1*, following band intensity quantification and adjustment for differences in amplicon size (non-adapted gel pictures depicted in Supplemental Figure S3B). B) Graph displaying change in *FN1*-EDA+ mRNA abundance over total *FN1* (*n* = 4). Lines connect biological replicates before and after differentiation. C) Western blot of FN1 and FN1-EDA in progenitor and differentiated keratinocytes. Protein abundances are shown as a percentage of progenitor state after blot quantification (non-adapted Western blot pictures depicted in Supplemental Figure S3C). Graph displaying the intensity of specific bands on Western blot, corrected for the housekeeping gene TUBB and normalized to the progenitor state (*n* = 2).

### *PRANCR* controls expression of splicing factors involved in EDA splicing

Splicing depends on interaction between SFs and their target pre-mRNAs (Barash *et al*, 2010). Expression levels of specific SFs are tightly regulated during tissue development and cell differentiation, which ensures expression of specific isoforms during different stages of tissue development (Baralle & Giudice, 2017). In the epidermis, regulated expression of SFs contributes to maintaining epidermal cells in an undifferentiated state (Tanis *et al*, 2018). We therefore aimed to identify the SFs that regulate *FN1*-EDA inclusion or exclusion in progenitor keratinocytes.

We first applied an *in silico* approach using the <u>d</u>ata<u>b</u>ase of <u>m</u>olecular <u>i</u>nteraction associated with <u>a</u>lternative <u>s</u>plicing (MiasDB), a tool that predicts interactions between mRNAs and SFs (Xing *et al*, 2016). MiasDB linked three members of the <u>s</u>erine/arginine-<u>r</u>ich (SR) family of <u>s</u>plicing <u>f</u>actors (SRSFs) to splicing of the FN1 EDA-exon: *SRSF1*, *SRSF3* and *SRSF7* (Figure 3A). All three SRSFs are highly expressed in progenitor keratinocytes compared to differentiated keratinocytes (Figure 3B), consistent with their having a role in controlling *FN1*-EDA splicing in progenitors. Splicing regulatory sequences for *FN1*-EDA are located within the exon itself (Mardon *et al*, 1987): a purine-rich <u>e</u>xonic <u>s</u>plicing <u>e</u>nhancer (ESE) region of 81 nucleotides that is recognized by SR proteins (Buratti *et al*, 2004). In hepatoma Hep3B cells, the ESE region enables binding of SRSF1 and SRSF7 leading to inclusion of the EDA exon in the *FN1* mRNA (Cramer *et al*, 1999; Lavigueur *et al*, 1993). Whether EDA is included in mature *FN1* mRNA is thought to depend on the interplay between SRSF1, SRSF7 and the physical properties of SRSF3 and the RNA polymerase II transcribing *FN1* (Cramer *et al*, 1999; Lavigueur *et al*, 1993; Buratti *et al*, 2004; Caputi *et al*, 1994; Kumar *et al*, 2019; Sen *et al*, 2015; Muro *et al*, 2008).

**Figure 3.**
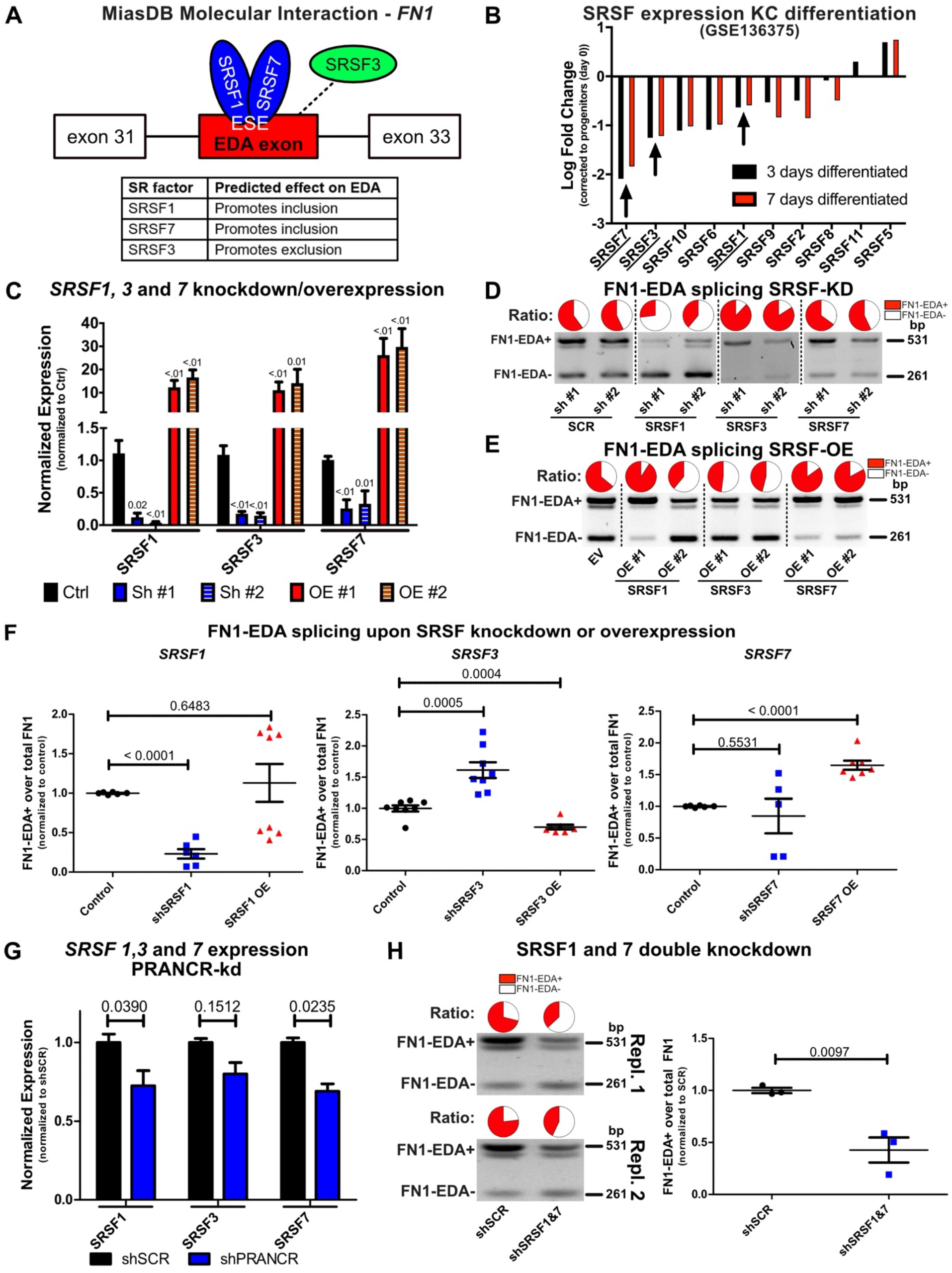
Regulation of FN1-EDA splicing by SRSFs and *PRANCR*. A) Schematic displaying the splicing factors indicated by MiasDB to regulate FN1-EDA splicing (Xing *et al*, 2016). B) Graph representing reanalysis of microarray data from progenitor and 3 or 7 days differentiated keratinocytes (GSE136375) using GEO2R. Bars represent Log2 Fold Change values of differentiated keratinocytes compared to progenitors (day 0) for all SRSF-family members present on the array (*n* = 4). C) mRNA expression of *SRSF1, 3,* and *7* in control (shSCR or Empty vector), *SRSF*-depleted progenitors (SRSF KD) and progenitors with *SRSF* overexpression (SRSF OE). Bars, mean with SEM. D) Representative RT-PCR gels showing differences in FN1-EDA abundance over total FN1 following band intensity quantification and adjustment for differences in amplicon size (non-adapted gel pictures depicted in supplemental Figure S4A). E) Representative gel showing the difference in *FN1*-EDA abundance in keratinocytes enriched for SRSF*1*, *3,* or *7* compared to cells transfected with an empty vector (EV), with ratios of *FN1*-EDA+ over total *FN1* calculated as in D (non-adapted gel pictures depicted in supplemental Figure S4B). F) Graphs showing calculated ratios of *FN1*-EDA+ over total *FN1*, normalized to control (SCR + EV). Each dot represents the outcome of an independent experiment. G) mRNA expression of *SRSF1*, *3*, and *7* in control and *PRANCR*-depleted cells in four independent biological replicates, retrieved from published RNA-seq data (GSE125400) (Cai *et al*, 2020). H) Left: representative gels of two replicate experiments showing differences in *FN1*-EDA+ abundance in keratinocytes depleted for both *SRSF1* and *SRSF7*, with ratios calculated as in D (non-adapted gel pictures depicted in supplemental Figure S4D). Right: Graph showing quantitation of *FN1*-EDA+ over total *FN1*, normalized to control (shSCR). Each dot represents outcomes of independent biological replicates.

To examine if these SRSFs control *FN1*-EDA+ splicing in keratinocytes, we depleted and overexpressed each SRSF in primary keratinocytes (Figure 3C) and used semi-quantitative RT-PCR to assess if these perturbations impacted the relative expression of the *FN1*-EDA+ isoform (Figure 3D-E and Supplemental Figure S4A-B). Depletion of *SRSF1* resulted in a decreased proportion of *FN1*-EDA+ (from ~0.60 in controls to 0.27-0.39 in SRSF1 knockdown), while overexpression did not result in clear significant changes (Figure 3D-F). For *SRSF7*, overexpression led to an increase in *FN1*-EDA+ (from 0.64 in controls to 0.83-0.85 in SRSF7 overexpression, Figure 3E), whereas depletion had no clear effect (Figure 3D-F). Lastly, for *SRSF3*, knockdown and overexpression led to opposite effects: its depletion led to an enrichment of *FN1*-EDA+ mRNA (from 0.61-0.64 to 0.84-0.88), whereas its overexpression led to decreased levels of FN1-EDA+ mRNA (from 0.64 to 0.46-0.48, Figure 3D-F). Together, these results demonstrated that alteration of SRSF1, 3, and 7 in primary keratinocytes are sufficient to influence FN1-EDA splicing.

We next assessed if *PRANCR* affected expression of these SRSFs. We observed decreased expression of *SRSF1* and *7* upon *PRANCR* depletion in RNA-seq data (Figure 3G), an observation confirmed with independent RT-qPCR experiments (Supplemental Figure S4C). *SRSF3* was also reduced but did not meet threshold for statistical significance (Figure 3G). Because *PRANCR* depletion led to a combined reduction of both SRSF1 and 7, we recapitulated this state using RNA interference (RNAi) targeting both *SRSF1* and *SRSF7* in wild-type keratinocytes. Dual knockdown led to a strong proportional reduction of *FN1*-EDA+ (from 0.71-77 to 0.37-0.43, Figure 3H and Supplemental Figure S4D). This result phenocopies the loss of *FN1*-EDA+ isoform predominance resulting from *PRANCR* depletion (Figure 1B-C) and indicated that the reduction of *SRSF1* and *SRSF7* that result from *PRANCR* depletion can recapitulate loss of FN1-EDA+ expression.

### *PRANCR* and *FN1*-EDA promote keratinocyte proliferation and migration

We next wanted to investigate how *FN1*-EDA+ expression affects keratinocyte function. Following skin injury and epidermal tissue loss, keratinocytes at the wound gap are mobilized to proliferate, migrate to fill the wound gap, and then stratify back into a multilayer tissue in a process known as re-epithelialization (Pastar *et al*, 2014). *FN1*-EDA+ expression increases at the base of the wound following injury (Ffrench-Constant *et al*, 1989). *FN1*-EDA-deficient mice show delayed reepithelization (Muro *et al*, 2003). These observations indicate a potential role for *FN1*-EDA+ and *PRANCR* in keratinocyte proliferation and migration, two key elements of epithelial wound healing.

We developed RNAi reagents to experimentally discriminate between total *FN1* and *FN1*-EDA+ isoforms. We depleted total *FN1* using RNAi targeted to universal *FN1* 3’ UTR sequences, while we depleted *FN1*-EDA+ isoforms by specifically targeting the EDA-containing exon (Figure 4A and Supplemental Figures S5A-B). To confirm experimental reduction of mRNA and protein *FN1* isoforms, we used semi-quantitative RT-PCR and Western blotting. RNAi targeting of the *FN1*-3’ UTR successfully depleted all *FN1* isoforms on both RNA (Figure 4B and Supplemental Figure S5C) and protein levels (Figure 4C-D and Supplemental Figure S5D). Specific targeting of the EDA-containing exon exclusively depleted *FN1*-EDA+ isoform expression (Figure 4B and Supplemental Figure S5C), resulting in an almost complete loss of the FN1-EDA+ protein isoform (<5% of control expression, Figure 4C-D and Supplemental Figure S5D). Furthermore, RNAi targeting of *FN1*-EDA led to only partial loss of total FN1 protein (~61% of control, Figure 4C-D, Supplemental Figures 5D-E), suggesting that in primary keratinocytes ~40% of total FN1-proteins contain the EDA domain.

**Figure 4.**
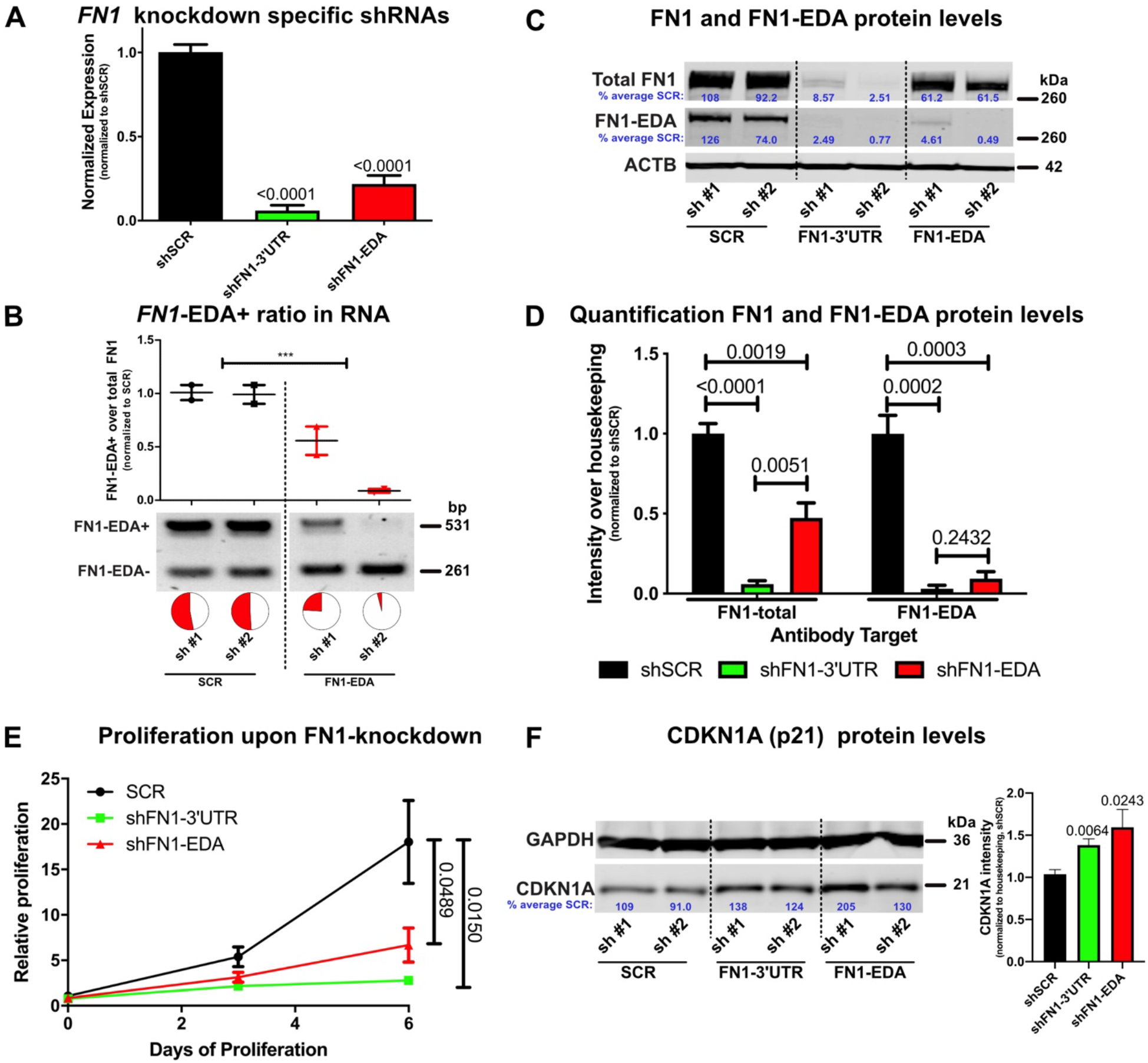
The FN1-EDA domain promotes keratinocyte proliferation. A) mRNA expression of total *FN1* in control (shSCR) and *FN1*-depleted cells after targeting total *FN1* in the 3’UTR region or by targeting the *FN1*-EDA exon (individual shRNA results in Supplemental Figure S5B). B) Representative gels showing differences in *FN1*-EDA abundance over total *FN1* following band intensity quantification and adjustment for differences in amplicon size (non-adapted gel pictures depicted in supplemental Figure S5C) in control (SCR) and *FN1*-EDA depleted (*FN1*-EDA) keratinocytes. C) Western blot of FN1 and FN1-EDA in control (SCR), *FN1*-depleted (FN1-3’UTR) and *FN1*-EDA-depleted (FN1-EDA) keratinocytes. Protein abundances are shown as a percentage of SCR after both quantification after normalization for total protein levels, as assessed by ACTB (non-adapted western blot pictures depicted in supplemental Figure S5D). D) Graph shows intensity of specific bands on Western blot, corrected for the housekeeping protein ACTB and normalized to controls (*n* = 4). E) Proliferation assay of control (SCR) and *FN1* (shFN1-3’UTR) or *FN1*-EDA (shFN1-EDA) depleted progenitors, measured with a fluorescence-based cell quantitation assay. Plotted values represent relative increase at each time point relative to day 0. n = 4; dots represent mean value with SEM. F) Western blot of CDKN1A in control (SCR), *FN1*-depleted (FN1-3’UTR) and *FN1*-EDA-depleted (*FN1*-EDA) keratinocytes. Protein abundances are shown as a percentage of SCR after both quantification after normalization for total protein levels, as assessed by GAPDH (non-adapted western blot pictures depicted in supplemental Figure S5G). Graph shows the intensity of CDKN1A bands on Western blot, corrected for the housekeeping protein GAPDH and normalized to controls (*n* = 4).

We assessed the proliferation rate of progenitor keratinocytes after genetic depletion of total *FN1* or *FN1*-EDA+. We observed that upon loss of total *FN1*, keratinocyte proliferation was severely inhibited (Figure 4E, Supplemental Figure 5F) and protein levels of CDKN1A, a potent inhibitor of cell cycle progression, were increased by >1.25x (Figure 4F), confirming a requirement for *FN1* in keratinocyte proliferation. Interestingly, RNAi targeting of the *FN1*-EDA domain, despite leaving significant residual FN1-EDA-negative protein (Figure 4D), also resulted in a major loss of keratinocyte proliferative capacity (Figure 4E) and increased CDKN1A protein levels (Figure 4F). This result suggested that the *FN1*-EDA+ isoform has a major role in maintaining keratinocyte proliferative capacity. Consistent with this, the RNAi targeting construct in which the FN1-EDA+ isoform was less completely depleted (shFN1-EDA #1, Figure 4B-C) showed greater residual proliferative rates (Supplemental Figure S4F) compared with keratinocytes in which the *FN1*-EDA+ isoform was more completely depleted (shFN1-EDA #2, Figure 4B-C). Others have observed that the addition of EDA peptide to culture medium of embryonic stem cells stimulates their proliferation (Losino *et al*, 2013) and that synchronized HaCaT keratinocytes have the highest level FN1-EDA+/EDA- ratio during active growth (Szell *et al*, 2004). Viewed together with our results, this indicates that the *FN1*-EDA+ isoform promotes keratinocyte proliferation.

Because *FN1* is part of the KCF network regulating both cell proliferation and cell mobility/adhesion-networks (Wu *et al*, 2012), we next turned our attention to whether *FN1*-EDA+ also influences the migrative capacity of keratinocytes. We seeded keratinocytes in chamber slides containing fixed-width inserts that, upon removal, produce standardized 500 μm cell-free gaps (Supplemental Figure S6A). Wild-type keratinocytes rapidly migrated into the gap, resulting in closure after ~9 hours (Figure 5A). Cells depleted for total *FN1* and *FN1*-EDA+ displayed markedly delayed gap closure (Figure 5A-B), suggesting that the reepithelization defect previously observed in FN1-EDA deficient mice (Muro *et al*, 2003) is at least partly due to a deficiency in both keratinocyte proliferation and migration. This observation is corroborated by the observation that during skin injury, the pattern of *FN1* splicing normally switches predominantly to the EDA+ isoform (Ffrench-Constant *et al*, 1989), further stressing an important role for *FN1*-EDA+ isoforms in tissue development and repair.

**Figure 5.**
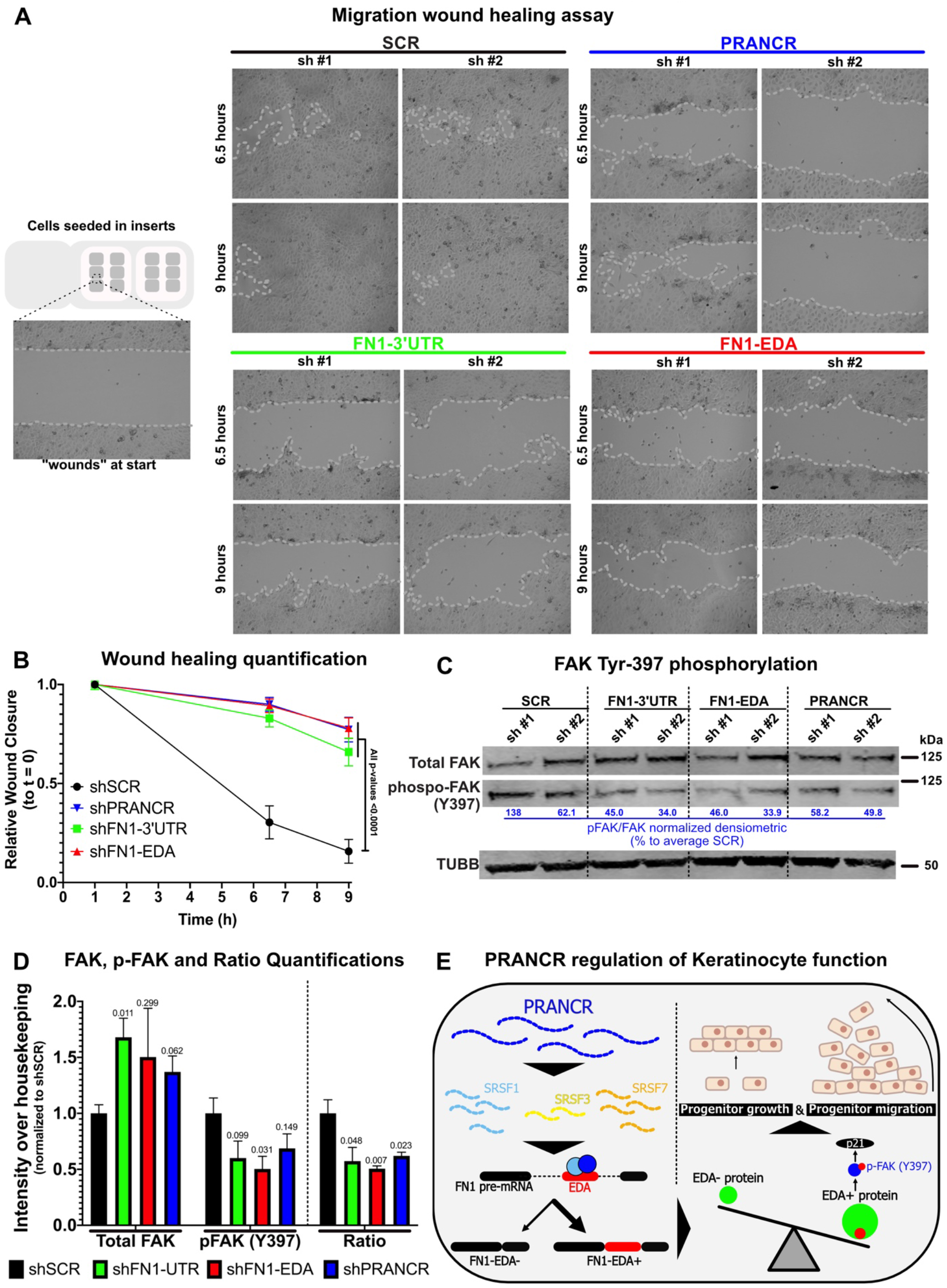
*PRANCR* and *FN1* are required for keratinocyte migration and FAK phosphorylation. A) Representative examples of images captured at 6.5 and 9 hours to display the process of *in vitro* wound gap closure with dotted lines indicating the boundaries of the wound. B) Graph indicating relative wound closure relative to the average wound sizes at the start, depicted as the area without cells (in area pixels^2) as assessed by the *Wound_healing_size_tool* ImageJ plugin (Suarez-Arnedo *et al*, 2020). Dots represent mean values with SEM (*n* = 12). C) Western blot of total FAK and FAK phosphorylated at Tyrosine-397 in control (SCR), *FN1*-depleted, *FN1*-EDA-depleted and *PRANCR*-depleted keratinocytes. Phosphorylated FAK over total FAK is shown as a percentage of the average value in control keratinocytes (SCR1+2; (non-adapted Western blot pictures depicted in Supplemental Figure S6B). D) Graph showing the intensity of Total FAK and phosphorylated FAK bands on Western Blots, corrected for the housekeeping protein TUBB and normalized to controls. Bars indicate average with SEM (*n* = 4). E) Working model how *PRANCR* affects keratinocyte behavior by regulating FN1-EDA splicing.

The FN1-EDA domain has been suggested to increase the binding affinity of fibronectin for integrins, the predominant FN1 receptor (Patten & Wang, 2021) Primary keratinocytes express 16 of the 26 integrin-encoding genes and many of them are required for keratinocyte function (Duperret *et al*, 2015). The FN1-EDA contains the EDGIHEL domain, which can bind to three different integrin receptors (Patten & Wang, 2021). In fibroblasts, signaling of FN1-EDA via these integrins leads to phosphorylation of focal adhesion kinase (FAK) at Tyrosine-397 (Kohan *et al*, 2010) and ensures LTBP1 protein levels (Klingberg *et al*, 2018). We therefore assessed whether key integrin signaling-related proteins are affected in keratinocytes depleted for either total *FN1* and *FN1*-EDA+. We observed decreased phosphorylation of FAK at Tyr-397 (Figure 5C-D and supplemental Figure 6B), while LTBP1 levels remained unchanged (Supplemental Figure 6C). As pFAK is required for skin formation (Duperret *et al*, 2015), keratinocyte migration and proliferation (Wang *et al*, 2018) and can regulate cellular levels of CDKN1A (Bryant *et al*, 2006), FN1-mediated phosphorylation of FAK provides a molecular mechanism by which *FN1*-EDA transduces its effect on keratinocyte cell fate.

Viewed together, these results lead us to propose a model in which *PRANCR* influences epidermal function, in part, by controlling the relative abundance of FN1-EDA+ through regulation of alternative splicing (Figure 5E). Given the involvement of both *PRANCR* and FN1-EDA+ in keratinocyte proliferation and migration, we hypothesize that the impact of these regulators on keratinocyte function may extend to key processes such as epidermal wound healing. Future studies will aim to explore the role of *PRANCR* in states of normal and impaired wound healing. The results of this work also highlight the broader need to understand the functional consequences of alternative splicing in the human genome. Up to 15% of disease-causing point mutations disrupt splicing (Krawczak *et al*, 1992), reflecting the involvement of AS in human disease. Understanding the interplay and functions of gene isoforms, splicing factors, and as well as the roles of lncRNAs in their regulation will contribute to our understanding of the functional complexity and diversity of the human genome.

